# National HIV testing and diagnosis coverage in sub-Saharan Africa: a new modeling tool for estimating the “first 90” from program and survey data

**DOI:** 10.1101/532010

**Authors:** M Maheu-Giroux, K Marsh, C Doyle, A Godin, C Lanièce Delaunay, LF Johnson, A Jahn, K Abo, F Mbofana, MC Boily, DL Buckeridge, C Hankins, JW Eaton

## Abstract

**Objective:** HIV testing services (HTS) are a crucial component of national HIV responses. Learning one’s HIV diagnosis is the entry point to accessing life-saving antiretroviral treatment and care. Recognizing the critical role of HTS, the *Joint United Nations Programme on HIV/AIDS* (UNAIDS) launched the 90-90-90 targets stipulating that by 2020, 90% of people living with HIV know their status, 90% of those who know their status receive antiretroviral therapy, and 90% of those on treatment have a suppressed viral load. Countries will need to regularly monitor progress on these three indicators. Estimating the proportion of people living with HIV who know their status (i.e., the “first 90”), however, is difficult.

**Methods:** We developed a mathematical model (henceforth referred to as “F90”) that formally synthesizes population-based survey and HTS program data to estimate HIV status awareness over time. The proposed model uses country-specific HIV epidemic parameters from the standard UNAIDS Spectrum model to produce outputs that are consistent with other national HIV estimates. The F90 model provides estimates of HIV testing history, diagnosis rates, and knowledge of HIV status by age and sex. We validate the F90 model using both in-sample comparisons and out-of-sample predictions using data from three countries: Côte d’Ivoire, Malawi, and Mozambique.

**Results:** In-sample comparisons suggest that the F90 model can accurately reproduce longitudinal sex-specific trends in HIV testing. Out-of-sample predictions of the fraction of PLHIV ever tested over a 4-to-6-year time horizon are also in good agreement with empirical survey estimates. Importantly, out-of-sample predictions of HIV knowledge are consistent (i.e., within 4% points) with those of the fully calibrated model in the three countries, when HTS program data are included. The F90 model’s predictions of knowledge of status are higher than available self-reported HIV awareness estimates, however, suggesting –in line with previous studies– that these self-reports could be affected by non-disclosure of HIV status awareness.

**Conclusion:** Knowledge of HIV status is a key indicator to monitor progress, identify bottlenecks, and target HIV responses. The F90 model can help countries track progress towards their “first 90” by leveraging surveys of HIV testing behaviors and annual HTS program data.

## Introduction

HIV testing services (HTS) are the entry point for diagnosis and access to life-saving antiretroviral therapy (ART)^[1]^. Early diagnosis and initiation of ART have been shown to drastically decrease viral load, which reduces individual morbidity and mortality, and limits onward HIV transmission^[2]^. HTS can also offer a pathway for primary prevention interventions, including programs that deliver pre-exposure prophylaxis, voluntary medical male circumcision, and prevention of mother-to-child-transmission.

Recognizing the critical role of HTS in a country’s national response, the *Joint United Nations Programme on HIV/AIDS* (UNAIDS) launched in 2014 the 90-90-90 targets stipulating that by 2020, 90% of people living with HIV (PLHIV) know their status, 90% of PLHIV who know their status receive ART, and 90% of those on treatment have a suppressed viral load^[3–5]^. To reach those targets, countries need to monitor progress on these three indicators, identify bottlenecks, and implement or adapt targeted testing and treatment services in a timely manner.

As of 2017, UNAIDS estimates that the biggest bottleneck globally in achieving the 90-90-90 targets is access to HIV testing, with about 25% of PLHIV estimated to not know their HIV status^[6]^. Estimating the proportion of PLHIV who know their status (i.e., the “first 90”), however, is difficult. In countries with robust and comprehensive HIV case surveillance systems, the proportion of diagnosed PLHIV can be estimated by triangulating HIV incidence and mortality with the cumulative number of new HIV diagnoses annually. In sub-Saharan Africa (SSA), where more than two-thirds of PLHIV reside^[7]^, surveillance systems are not sufficiently developed. Many countries estimate the proportion of PLHIV who know their status primarily from nationally representative household surveys.

Most *Demographic and Health Surveys* (DHS) and *AIDS Indicator Surveys* (AIS) in SSA include HIV serology, with respondents self-reporting whether they have ever been tested for HIV, but are rarely being asked directly if they are aware of their HIV status. The proportion of HIV positive respondents who report ever having been tested for HIV serves as an upper bound for the level of HIV awareness, since the last HIV test might have been HIV-negative (i.e., occurring before the person seroconverted). In recent years, *Population-based HIV Impact Assessments* (PHIA) surveys and a few other surveys conducted in SSA countries have collected information on both HIV seroprevalence and self-reported awareness status. These data have been used directly to estimate the “first 90”^[8, 9]^. However, comparison of self-reported awareness of HIV status with biomarker measurements of antiretroviral usage and viral load suppression reveals sometimes substantial non-disclosure of awareness of HIV status for persons who are on ART^[9, 10]^.

The infrequency of large population-based seroprevalence surveys, which are typically conducted every 5 years, also hampers regular monitoring of HIV awareness^[11]^. UNAIDS has previously estimated the change in knowledge of status over time in countries with survey data by applying additional increases in knowledge of status proportional to the scale-up in ART coverage between the current reporting year and the year of the last survey^[12]^. However, there is a need to better estimate progress towards the “first 90” in relation to changes in ART coverage and HTS program efforts^[13]^. For example, the relationship between ART coverage and knowledge of status has likely changed as a function of eligibility for treatment initiation. Further, programmatic data of the numbers of people tested and those testing HIV-positive could help to inform changes in testing levels.

To address these challenges, we developed a mathematical model –henceforth referred to as the “F90” model– that formally synthesizes population-based surveys and HTS program data within a Bayesian framework to estimate knowledge of status among people (≥15 years) over time in SSA. F90 estimates HIV testing and diagnosis rates over time by age, sex, and previous HIV testing history, to generate estimates of the “first 90” and other indicators of interest such as positivity among HIV testers and yield of new HIV diagnoses. Key features of this new F90 model are that it:

- takes as inputs –and therefore is fully consistent with– national modeled estimates of HIV prevalence, incidence, mortality, and ART coverage derived using the UNAIDS-supported Spectrum modeling software.
- uses data about self-reported HIV testing history from surveys that both include and do not include HIV serology.
- incorporates programmatic data on the annual numbers of HIV tests administered (e.g., testing volume) and number of positive HIV tests (positivity), where available.

## METHODS

### Modeling framework

HIV testing uptake is dynamically modeled using a deterministic framework based on a system of ordinary differential equations adapted from a well-established South African model^[14]^. The F90 model stratifies a country/region’s population by HIV testing history, and among PLHIV, by knowledge of HIV status among PLHIV, and ART status resulting in the following six main cascade stage: 1) HIV-susceptible who have never been tested, 2) HIV-susceptible ever tested, 3) PLHIV who have never been tested, 4) PLHIV unaware who have ever been tested, 5) PLHIV aware (untreated), and 6) PLHIV on ART. A schematic of the compartmental flows between these different stages is presented in Figure 1.

**Figure 1.**
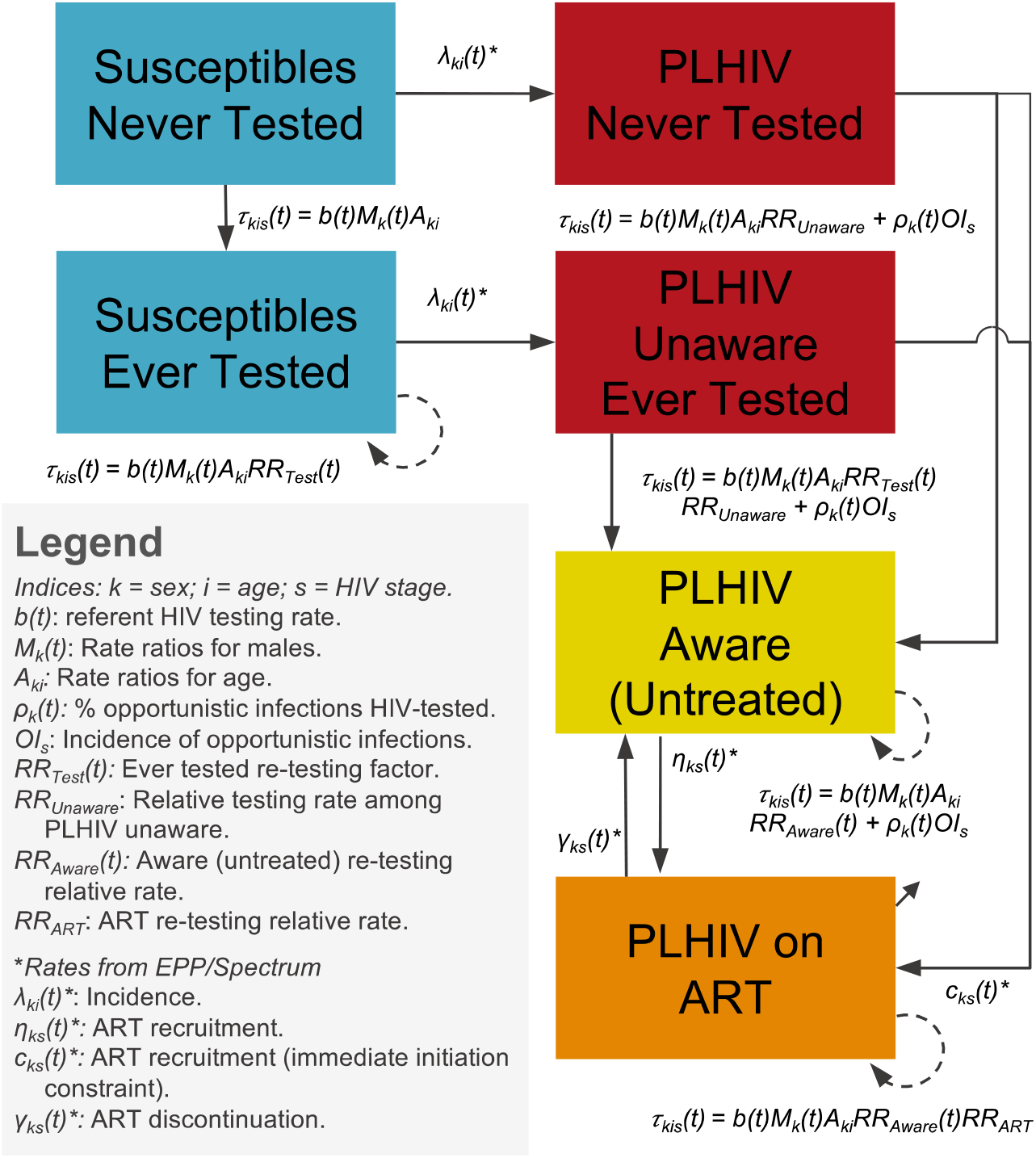
Inter-compartmental flow describing HIV testing uptake as a function of HIV status (susceptible versus living with HIV), testing history (never versus ever tested), HIV awareness status and antiretroviral treatment (ART) status. All parameters related to HIV testing (*τ*_*kis*_(*t*)) are estimated by the F90 model. Other rates, such as HIV incidence, ART recruitment and discontinuation are informed by Spectrum/EPP, as well as demographic parameters governing entry in the model at 15 years of age and both natural and HIV-related mortality (not depicted on the figure for ease of visual interpretation).

Individuals enter the population at age 15 years and are assumed to have never been tested for HIV (unless already living with HIV and on ART). F90 has been developed to use as inputs annual estimates of HIV incidence, mortality, and ART coverage produced by countries and published annually by UNAIDS^[15]^. At the core of the estimation process is Spectrum’s *AIDS Impact Module* and its *Estimation and Projection Package* (EPP)^[16]^. The Spectrum model, its assumptions, data requirements, and software are described in detail elsewhere^[17]^. Importantly for the new F90 model, Spectrum produces epidemic statistics stratified by age and sex, CD4+ cell count category, and ART status.

The transitions rates between HIV-susceptibles and PLHIV are informed by point estimates of the sex- and age-specific incidence rates estimated by Spectrum/EPP, as well as the transition rates from the three PLHIV untreated stages to the PLHIV on ART one (and ART discontinuation)^[17]^. Spectrum/EPP also informs demographic rates and HIV disease progression and mortality^[17]^. The F90 model is used to estimate all HIV testing rates, as further described in the next section.

### Model specification for HIV testing

The per capita rate *τ*_*kius*_(*t*) at which individuals are tested for HIV varies by calendar time (*t*), sex (*k*), age (*i*), HIV testing history and awareness status (*u*), and, for PLHIV, CD4+ cell count (*s*). Specifically, it takes the following form:

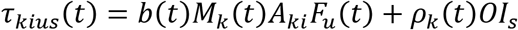

Here, *b(t)* is the testing rate for the referent group of women in the 15-24 years age category for calendar year *t*, which is assumed to be negligible in sub-Saharan Africa before 1995 (see Text S1 for full details). From 2000 onwards, *b(t)* is modeled as a first-order random walk with annual steps. *M*_*k*_(*t*) represents the HIV testing rate ratio for men (*k*=1) aged 15-24 years relative to women aged 15-24 at time *t* (equal to 1 for this referent group). We allow for changes in this ratio from 2005 and 2010 to account for potential scaling-up of prevention of mother-to-child transmission programs in sub-Saharan Africa countries^[18, 19]^, which could have influenced sex differences in HIV testing uptake. The term *A*_*ki*_ is the age- and sex-specific HIV testing rate ratio for ages 15-24 (*i*=1), 25-34 (*i*=2), 34-49 (*i*=3), and 50+ (*i*=4) age groups, which are assumed to be time-invariant^[8, 14, 20, 21]^.

The term *F*_*u*_(*t*) allows for potential differences in HIV testing rates according to prior HIV testing history and HIV status between HIV-susceptible who have never been tested (*u*=1), HIV-susceptible previously tested (*u*=2), PLHIV who have never been tested (*u*=3), PLHIV unaware who have ever been tested (*u*=4), PLHIV aware not on treatment (*u*=5), and PLHIV on ART (*u*=6); as displayed in Figure 1 and further described below.

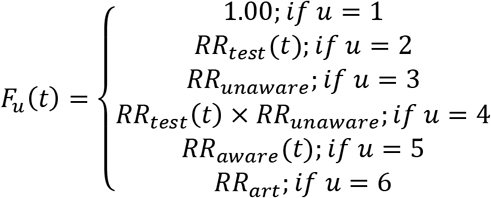

In many SSA countries, a substantial fraction of the population is tested every year but the proportion of people reporting being ever tested remains lower than what would be expected if everybody in the population had tested at an equal rate. Empirical evidence also suggests that rates of HIV testing are higher among people who have previously been tested for HIV^[22–26]^. The *RR*_*test*_(*t*) rate ratio is introduced to take this into account. PLHIV who are unaware of their status could also test at higher or lower rates than those who are HIV-susceptible individuals. Hence, differential testing rates in this group is accounted for with the *RR*_*unaware*_ rate ratio. Further, the number of positive tests is often very large, such that the cumulative number of positive HIV tests reported by HTS programs substantially outstrips the number of PLHIV who could have been newly diagnosed. This suggest that a non-negligible fraction of PLHIV aware of their status and PLHIV receiving ART may also be re-tested for HIV each year^[27, 28]^. For example, in many countries (e.g., Côte d’Ivoire^[29]^; Mozambique, F. Mbofana, pers. comm.; Senegal^[30]^; Sierra Leone^[31]^; Uganda^[32]^), the annual numbers of positive tests reported often represent up to 25-30% of the whole estimated PLHIV population, which is inconsistent with survey data on the proportion of PLHIV ever tested. To reproduce the number of positive tests, and in line with empirical evidence, we allowed re-testing of diagnosed PLHIV using the time-varying *RR*_*aware*_(*t*) rate ratio. Finally, PLHIV on treatment could also be retested for HIV, albeit at lower rate through the *RR*_*art*_ rate ratio. The main studies informing differential testing rates are summarized in supplementary material (Tables S2-S3).

Lastly, we consider that HTS uptake will depend on the proportion of untreated PLHIV experiencing HIV/AIDS-related symptoms who are not on ART. *OI*_*s*_ is the time-invariant incidence of opportunistic infection by CD4+ cell count category *s* ^[33, 34]^ (as tracked in Spectrum) and *ρ_k_(t)* is the sex-specific proportion of these infections that are tested for HIV at time *t* (see Text S1 for full details on the F90 model).

### Data sources, likelihood function, and model calibration

Two main data sources are used for model calibration: 1) household survey data about the proportion of adults who self-report having ever been tested for HIV and 2) HTS program data about the total number of HIV tests conducted each year and number of HIV positive tests. For national surveys, we used the proportion of respondents reporting having “*ever been tested and received the result of the last HIV test*”, stratified by sex, age (15-24, 25-34, and 35-49), and, if available, HIV serostatus from nationally representative household surveys, including DHS, AIS, *Multiple Indicator Cluster Surveys* (MICS), PHIA surveys, and relevant country-specific surveys (e.g., HSRC, KAIS). For surveys that do not include HIV serostatus (e.g., some MICS and DHS), data about ever testing by age and sex, irrespective of HIV status, are used in model calibrations. We assume that self-reports of “*ever having been tested and receiving the result of the last HIV test*” are unbiased estimates of HIV testing history.

For HTS program data, the F90 model can also be calibrated to the annual number of HIV tests performed in the population (≥15 years) and, if available, the number of positive tests (≥15 years; stratified by sex or overall). Such data may be useful to inform testing trends after the last population-based surveys have been performed. Details of the likelihood specification can be found in the supplemental material (Text S2).

Notably, we purposely excluded two commonly referenced data types from surveys as potential inputs into F90. First, information on HIV testing in the past year is not used in model calibration due to evidence that this likely overstates the true annual testing rate^[14, 35]^, perhaps due to “telescoping bias” in which respondents may inadvertently recall testing that occurred beyond the last 12 months^[36]^ (see supplemental materials, Text S3). Second, information on self-reported awareness of HIV-positive status, even when partially adjusted for detection of ART among PLHIV who report not knowing their status, is not incorporated due to evidence of systematic nondisclosure of knowledge of status^[9, 10, 37–44]^. In particular, non-disclosure of HIV status was found to be 1.4 times higher among individuals not on ART in Mozambique^[38]^ compared to those on ART. This implies that adjustments for presence of ART metabolites may be insufficient, especially when ART coverage is low.

Model parameters were estimated using a Bayesian framework. To constrain the parameters space to plausible values in data-limited settings, we elicited prior distributions following a review of the literature (Tables S1-S2; prior distributions are described in Text S1). Posterior modes of model parameters were obtained via non-linear optimization using the Broyden-Fletcher-Goldfarb-Shanno algorithm^[45]^. The joint posterior distribution was estimated using a Laplace approximation^[46, 47]^, and 95% credible intervals for quantities of interest were obtained by sampling 3,000 parameter sets from this approximated joint posterior distribution and summarizing the posterior of relevant outputs using the median, and 2.5^th^ and 97.5^th^ percentiles. This calibration method was chosen for its computational efficiency. Table S3 presents comparisons of summary statistics of the posterior distributions of selected model outputs using the Laplace approximation, *Sampling Importance Resampling* (SIR), and the *Incremental Mixture Importance Sampling* (IMIS)^[48]^ algorithms. These suggest good performance of the Laplace approximation in our settings.

All analyses were conducted in the R statistical software^[49]^. The system of ordinary differential equations was solved using a Euler algorithm with a time step of 0.1 years. All functions are available for download from a Github repository (https://github.com/mrc-ide/first90release).

### Model outputs (estimates)

The F90 model generates results for comparisons to input data and indicators of interest. It estimates the total number of tests (negatives and positives), tests among first-time testers, positivity (the percent of positive tests among all tests) and yield (the percent of new HIV diagnoses among all tests), the proportion of the population ever tested for HIV, the proportion of PLHIV who know their HIV status, and other indicators (Table S4).

### Model validation

There are few empirical estimates of awareness status among PLHIV and, as described above, these estimates are likely to reflect substantial underreporting of HIV awareness^[9, 10, 37–44]^. We therefore validated the F90 model by performing both in-sample comparisons (A) and out-of-sample predictions (B and C) of the proportion of the population ever tested for HIV (stratified by sex and HIV status). We focused our analyses on three countries with multiple surveys and availability of HTS program data: Côte d’Ivoire, Malawi, and Mozambique. For the out-of-sample predictions, we first excluded all surveys conducted after 2012 and all HTS program data after the last available pre-2012 survey (B). This was performed to examine the F90 model’s ability to predict testing histories over a time horizon of approximately 5 years (the time interval often observed between two population-based surveys). We then re-calibrated the model, this time incorporating the post-2012 HTS program data. To appreciate the added value of the HTS program data sources, we re-calibrated our model both on the sex-combined (C1) and sex-disaggregated HTS data (C2). In the case of Mozambique, available HTS program data were not stratified by sex and we instead used the fully-calibrated model (A) to predict sex-stratified HTS program data (2009-2017) which was then used for the out-of-sample validation (C2). The data sources used for model calibration are presented in Table 1.

**Table 1.** List of surveys of with information on the proportion of respondents having ever been tested for HIV (2000-2017) and HIV testing services program data used to calibrate the F90 model in Côte d’Ivoire, Malawi, and Mozambique.

| Data Types | Côte d’Ivoire | Malawi | Mozambique |
| --- | --- | --- | --- |
| **Surveys** | MICS 2000* (women only) | DHS 2004 | DHS 2003 |
|  | AIS 2005 | MICS 2006* (women only) | MICS 2008* (women only) |
|  | DHS 2012 | DHS 2010 | AIS 2009 |
|  | MICS 2016* (E) | MICS 2014* (E) | DHS 2011* |
|  | PHIA 2017† (E) | DHS 2015 (E) | AIS 2015 (E) |
|  |  | PHIA 2016 (E) |  |
| **HIV testing services program data** | *Direction de l’information, de la planification et de l’évaluation.* (total tests and number positives for 2010-17; sex-disaggregated for 2014-17) | *Malawi Integrated HIV Program Report.* (total tests and number positives 2003-17; sex-disaggregated for 2013-17‡) | *National HIV/AIDS Control Program (F. Mbofana, personal communication).* (2013-2017; total tests and number positives) |

AIS: AIDS Indicator Survey; DHS: Demographic and Health Survey; MICS: Multiple Indicator Cluster Survey; PHIA: Population-based HIV Impact Assessment.
(E): Indicates that the survey was excluded in the out-of-sample validation analyses.
\*Survey does not include serology and estimates of “ever tested for HIV” cannot be stratified by HIV status.
†The 2017 PHIA survey in Côte d’Ivoire has yet to be released in the public domain. Preliminary results for this survey were used in the model calibration and validation but the relevant point estimates cannot be presented at this time.
‡Only the total number of sex-disaggregated tests is available and the number of positive tests is for both sex combined.

### Ethics

All analyses were performed on anonymized and de-identified data. Further, all DHS/AIS survey protocols have been approved by the *Internal Review Board* of ICF International in Calverton (USA) and by the relevant country authorities for other surveys (MICS and PHIA). Further information on the ethics approval can be found in the individual country reports. Ethics approval was obtained from McGill University’s Faculty of Medicine *Institutional Review Board* (A10-E72-17B).

## Results

### Description of survey data on HIV testing history and HTS programs data

In each of the three countries, the survey-estimates for the proportion of the population reporting having ever been tested for HIV and receiving the last test’s result increased from under 15% at the beginning of the 2000s to 50% in Côte d’Ivoire (2016), 75% in Malawi (2016), and 51% in Mozambique (2015) (Figure 2). Women are more likely to report having ever been tested than men. As for age, the highest proportions of participants reporting a history of HIV testing is consistently found in the 25-34-year-old age group in all three countries. Testing among PLHIV is higher than in the general population, with survey estimates indicating that 68% and 93% of PLHIV Malawi (in 2016) and Mozambique (in 2015), respectively, report a history of HIV testing (Figure 2).

**Figure 2.**
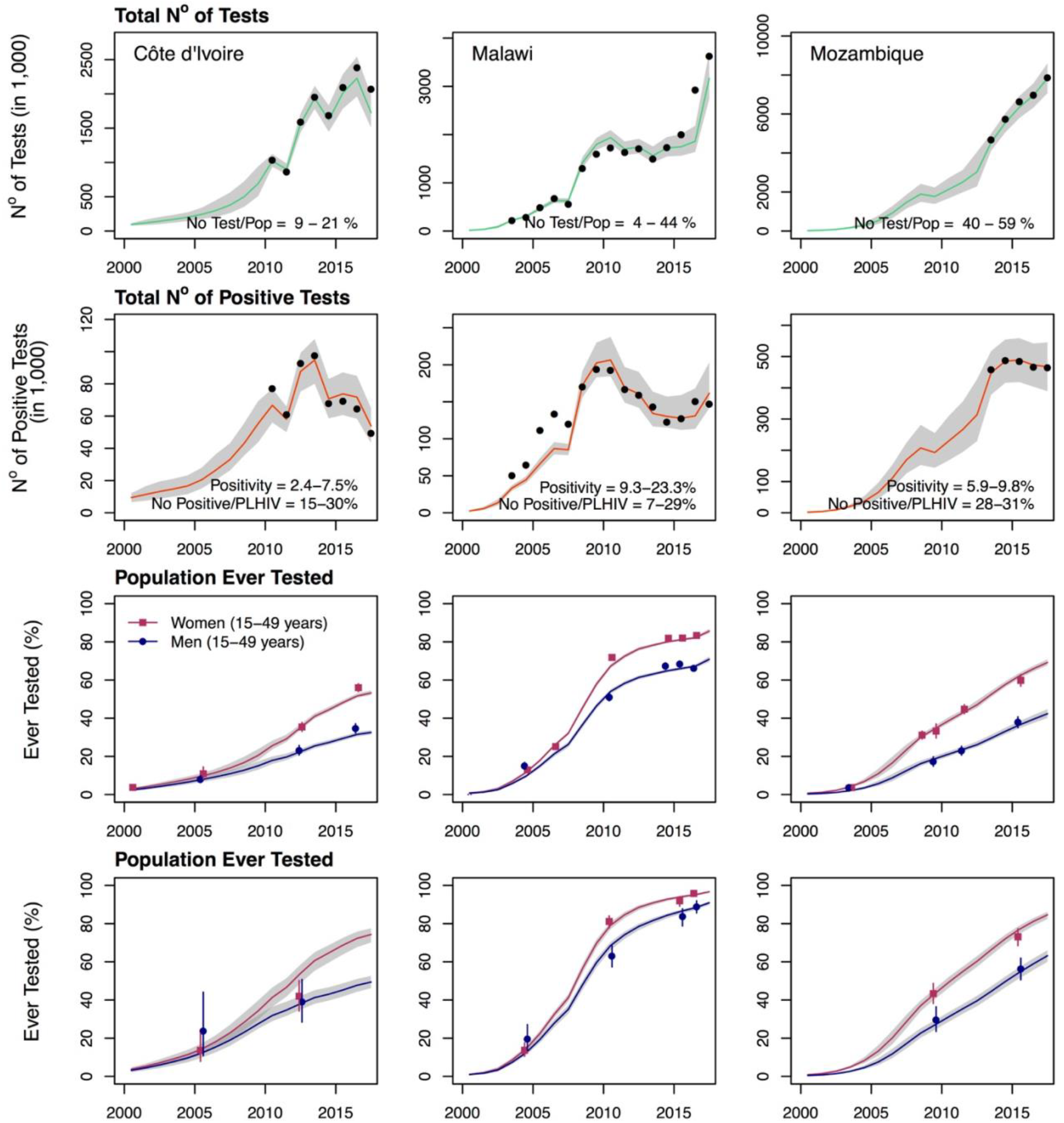
Comparison of calibrated F90 model fits with programmatic and survey data for Côte d’Ivoire (1^st^ column), Malawi (2^nd^ column), and Mozambique (3^rd^ column) over 2000-2017. The shaded areas on all graphs correspond to the 95% credible intervals of the posterior estimates, with the lines corresponding to the median. Black dots on the first and second row of graphs correspond to the reported HIV testing services program data for the overall number of tests (top row) and number of positive tests (2^nd^ row; see Table 1 for details). The points on the 3^rd^ and 4^th^ rows of graphs are the survey estimates of the proportion ever tested among women (squares) and men (circles) among the overall population (3^rd^ row) and people living with HIV (PLHIV; bottom row). The lines crossing the points are the 95% confidence intervals of the survey estimates.

HTS program data suggest that a substantial number of HIV tests are administered annually. For example, the reported maximum annual number of tests performed corresponds to 21% of the population aged 15-49 years old in Côte d’Ivoire, 49% in Malawi, and 59% in Mozambique (Figure 2). Concomitant with important increases in total testing volume, the number of positive tests has decreased in all three countries, resulting in downward trends in positivity rates. In addition, the number of positive tests reported in HTS program data suggests that a substantial fraction of diagnosed PLHIV could be retested every year. For example, the volume of positive tests corresponds to the equivalent of up to 30% of the total PLHIV population aged 15-49 years in Côte d’Ivoire, 29% in Malawi, and 31% in Mozambique. If these were all new diagnoses, we would expect that close to all PLHIV should be aware of their status within a few years.

#### A) In-sample comparisons – calibration on all available survey and HTS program data

The calibrated F90 models for Côte d’Ivoire, Malawi, and Mozambique can accurately reproduce annual HTS program data both for the total number of HIV tests performed and the number of positive tests (Figure 2). In addition, the model adequately reflects sex-specific survey estimates of the proportion of respondents ever tested for HIV. In 2017, these were estimated by F90 to be of 33%, 71%, and 42% among men in Côte d’Ivoire, Malawi, and Mozambique, respectively. Testing was notably higher among women, with 53% (Côte d’Ivoire), 86% (Malawi), and 69% (Mozambique) of women reporting having ever been tested for HIV. Overall for 2017, average testing rates were estimated to be 0.17 per year in Côte d’Ivoire, 0.21 per year in Malawi, and 0.52% per year in Mozambique.

The F90 model is also able to replicate longitudinal trends in the proportion of PLHIV ever tested. It estimates that 66%, 95%, and 76% of PLHIV in Côte d’Ivoire, Malawi, and Mozambique, respectively, have ever been tested for HIV in 2017. In turn, knowledge of HIV status is estimated by F90 at 58% in Côte d’Ivoire, 84% in Malawi, and 72% in Mozambique. These numbers are within the range of values obtained from the previous UNAIDS methodology for 2017 in Côte d’Ivoire (54%; uncertainty range 38-75%) and Malawi (90%; 84 to >95%), but the previous estimate was lower in Mozambique (59%; 49-70%)^[6]^.

F90 also suggests important differences in knowledge of status by sex, with higher proportions of women being aware of their status than men. HIV status knowledge is greatest in older age groups in all three countries. It increases from 41% among 15-24-year-olds to 65% among 35-49-years-olds in Côte d’Ivoire, from 69% (15-24 years) to 89% (35-49 years) in Malawi, and from 57% (15-24 years) to 74% (35-49 years) in Mozambique for the year 2017. As expected, the proportions of PLHIV aware of their status are higher than survey estimates of self-reported awareness, when these are available (Table 2 and Figure 3), consistent with previous literature suggesting non-disclosure of awareness of HIV status. In Malawi, estimates of knowledge of status are generally between estimates of PLHIV ever tested and the proportion on ART. In Côte d’Ivoire and Mozambique, the “first 90” is closer to the proportion of PLHIV ever tested because survey and HTS program data suggest high rates of re-testing. For example, the rate ratios for re-testing were estimated to be 3.7 (95% credible interval [95%CrI]: 3.2-4.3) in Côte d’Ivoire and 7.2 (95%CrI: 6.4-7.7) in Mozambique, as compared to 1.2 (95%CrI: 1.1-1.4) in Malawi. Posterior estimates for the main model parameters are reported in supplemental materials (Table S5).

**Table 2.** Comparisons of empirical survey estimates of the proportion of individuals aged 15-49 years old ever tested for HIV (by sex and HIV status) and self-reported awareness status among PLHIV with F90 model predictions from A) the fully calibrated model and from out-of-sample predictions that B) excluded all post-2012 survey and HIV testing services (HTS) program data, C1) excluded all post-2012 survey data (included sex-combined HTS program data), and C2) excluded all post-2012 survey data but included sex-disaggregated HTS data.

| Country / Outcome | Comparisons |  | Predictions (95% CrI) |  |  |  |
| --- | --- | --- | --- | --- | --- | --- |
|  | Survey & Year | Survey estimates (95% CI) | A) Full data calibration | B) Excluding survey and HTS data | C1) Excluding survey data only (HTS data sex-combined) | C2) Excluding survey data only (HTS data sex-disaggregated) |
| **Côte d’Ivoire** |  |  |  |  |  |  |
| Women ever tested | MICS 2016 | 56% (54-58%) | 52% (51-53%) | 51% (43-65%) | 53% (49-58%) | 53% (51-55%) |
| Men ever tested | MICS 2016 | 35% (32-37%) | 32% (30-33%) | 35% (28-46%) | 36% (31-41%) | 29% (28-31%) |
| WLHIV ever tested | PHIA 2017 | NPD | 74% (70-77%) | 72% (64-83%) | 74% (69-79%) | 74% (73-76%) |
| MLHIV ever tested | PHIA 2017 | NPD | 49% (46-52%) | 52% (44-66%) | 54% (48-61%) | 47% (45-49%) |
| PLHIV aware (“first 90”) | PHIA 2017 | 37%* | 58% (53-61%) | 56% (47-70%) | 59% (53-64%) | 57% (55-58%) |
| **Malawi** |  |  |  |  |  |  |
| Women ever tested | PHIA 2016 | 83% (82-84%) | 82% (81-83%) | 88% (77-96%) | 91% (90-92%) | 89% (87-90%) |
| Men ever tested | PHIA 2016 | 66% (65-68%) | 67% (66-68%) | 61% (51-76%) | 64% (60-68%) | 68% (66-70%) |
| WLHIV ever tested | PHIA 2016 | 96% (94-97%) | 95% (94-96%) | 98% (93-100%) | 98% (98-99%) | 98% (97-98%) |
| MLHIV ever tested | PHIA 2016 | 89% (86-92%) | 88% (87-89%) | 86% (79-93%) | 87% (84-89%) | 90% (88-91%) |
| PLHIV aware (“first 90”) | PHIA 2016 | 76% (73-78%) | 81% (79-82%) | 81% (73-89%) | 83% (82-85%) | 84% (82-85%) |
| **Mozambique** |  |  |  |  |  |  |
| Women ever tested | AIS 2015 | 60% (57-63%) | 62% (61-64%) | 56% (48-67%) | 70% (64-77%) | 65% (62-68%) |
| Men ever tested | AIS 2015 | 38% (35-41%) | 36% (35-38%) | 28% (24-37%) | 38% (32-46%) | 38% (36-40%) |
| WLHIV ever tested | AIS 2015 | 73% (68-77%) | 77% (75-79%) | 74% (67-83%) | 84% (78-89%) | 80% (76-83%) |
| MLHIV ever tested | AIS 2015 | 56% (51-62%) | 55% (51-57%) | 48% (42-56%) | 57% (50-64%) | 57% (53-60%) |
| PLHIV aware (“first 90”) | AIS 2015 | 40% (37-44%) | 63% (60-65%) | 52% (44-63%) | 68% (61-72%) | 66% (61-69%) |

95%CI: 95% confidence interval; 95% CrI: 95% credible interval; HTS: HIV testing services; NPD: not in public domain (but included in the calibration); MICS: multiple indicators cluster survey; MLHIV: men living with HIV; NA: not available; PHIA: population-based HIV impact assessment; WLHIV: women living with HIV.
\*Age group is 15-64 and estimate is not adjusted for antiretroviral metabolites.

**Figure 3.**
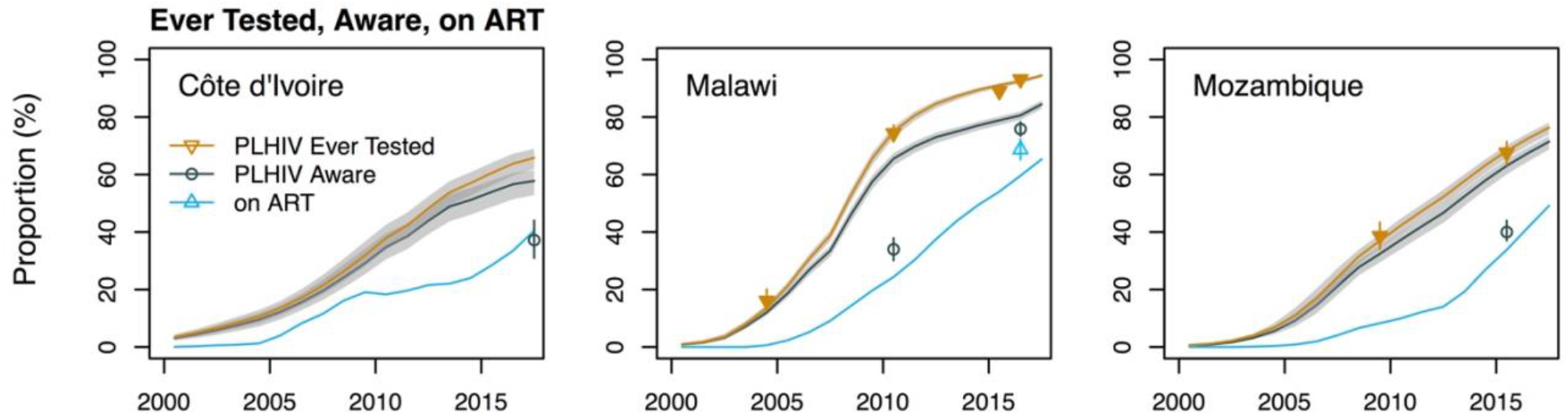
Comparisons of calibrated F90 model fits with survey data on proportion of people living with HIV (PLHIV) aged 15-49 years ever tested, model-predicted proportion of PLHIV aware of their status (“first 90”), and survey estimates of awareness status and Spectrum/EPP’s antiretroviral therapy (ART) coverage estimates. The shaded areas correspond to the 95% credible intervals of the posterior estimates. Estimates used for cross-validation are shown as empty symbols. (The self-reported estimate of awareness in Côte d’Ivoire correspond to the 15-64 age group.)

#### B) Out-of-sample predictions: removing both recent survey and HTS program data

When excluding all data from surveys conducted after 2012 and HTS program data after the last survey, the F90 model predictions underestimate by 6% points the 2016 survey estimate of the proportion of the population ever tested for HIV in Côte d’Ivoire (susceptible and PLHIV combined). In Malawi, the model’s out-of-sample predictions for 2016 are higher than the survey estimates among women by 5% points and underestimate the same proportion among men by 5% points, but both are included within the predicted uncertainty intervals (Table 2 and Figure 4). Finally, in Mozambique the proportion of women ever tested is overestimated by 4% in 2015 but, for men, the underestimation is 10% points for that same year.

**Figure 4.**
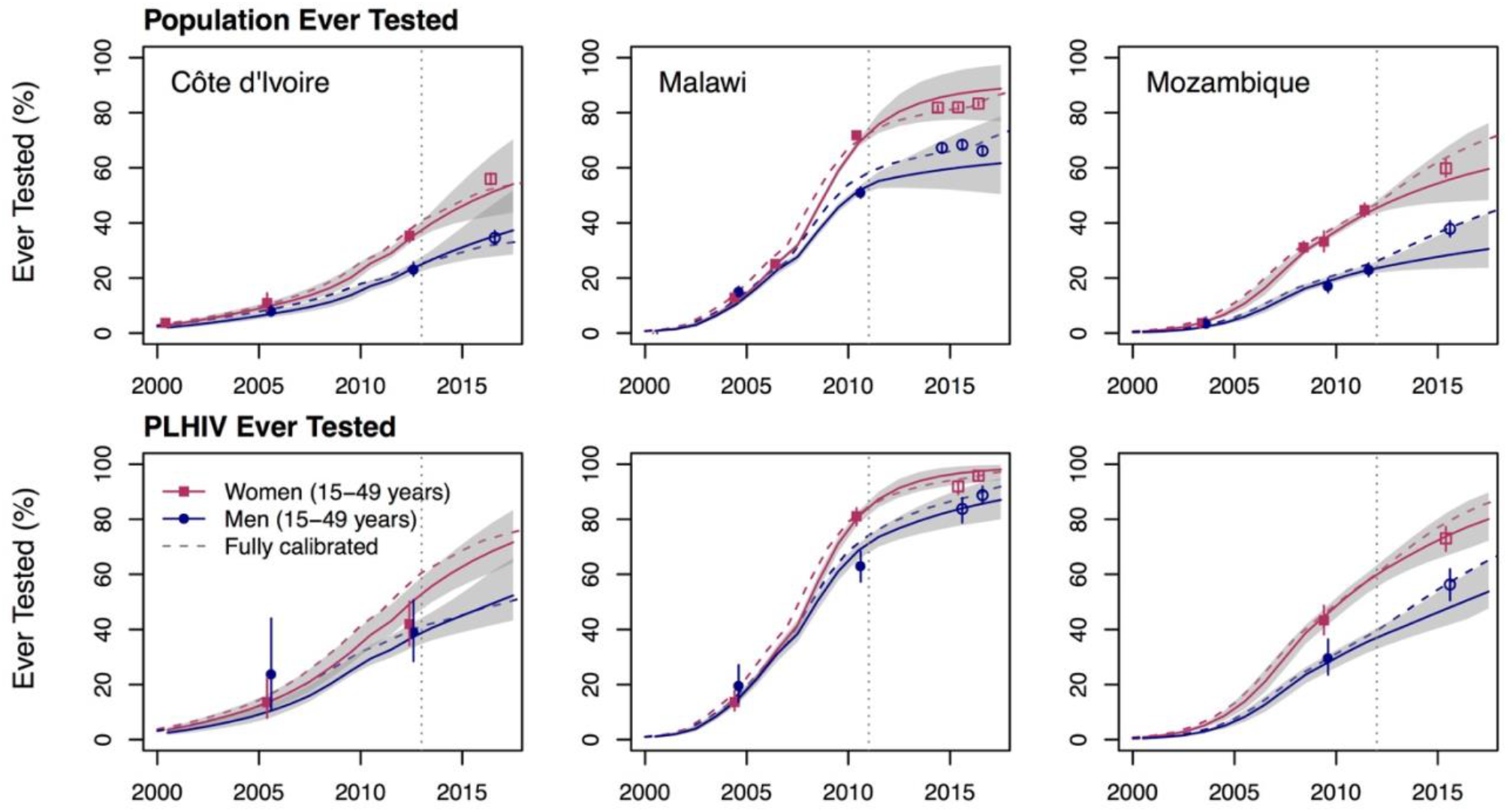
Out-of-sample predictions (B; full lines) of F90 models calibrated to survey data from 2000-2012, excluding all program data, for Côte d’Ivoire, Malawi, and Mozambique and model predictions for the 2013-2017 period. Dashed lines represent predictions from the fully calibrated models. The vertical lines indicate the date of the last survey data estimates included in the fitting (to the right of the lines are the predictions). The shaded areas correspond to the 95% credible intervals of the posterior estimates. Points represent the survey estimates (Table 1) and empty symbols indicate that these survey outcomes were not included in the likelihood but are shown for cross-validation purposes. The vertical solid lines crossing these points correspond to the 95% confidence intervals of the survey estimates. (PLHIV: people living with HIV.)

Estimates of ever testing among PLHIV are, arguably, a more relevant outcome to the “first 90” than corresponding estimates among the overall population. For PLHIV, out-of-sample predictions are quite accurate, even over a full 5-year time horizon for the three countries (Figure 4). In Côte d’Ivoire, the difference between the 2017 model prediction of the fraction of PLHIV ever tested and the empirical estimates is less than 3% points (for both sexes combined; empirical estimates from the Côte d’Ivoire PHIA are not yet in public domain - not shown). A similar pattern is observed for Malawi with differences of less than 1% and 3% points between predictions and empirical estimates for women and men, respectively. In Mozambique, there are differences of 4% and 10% points of the proportion ever tested among men and women, respectively. As for the proportion of PLHIV aware of their status in 2017, the out-of-sample predictions are within 2% points of the ones obtained using full data calibration in Côte d’Ivoire and Malawi. In Mozambique, the difference is 13% points but the uncertainty intervals are very wide and encompass the estimate from the full data calibration.

#### C) Out-of-sample predictions: added value of HTS program data

Adding the post-2012 HTS program data (sex-combined) yields mixed results with respect to improving estimates (Figure 5, C1). It adds little to the already accurate predictions in Côte d’Ivoire. In Malawi, however, it improves estimates for women but magnifies the underestimation in men (Table 2, Figure 5). For Mozambique, HTS program data increase the accuracy of predictions for men but result in overestimating the proportion of women ever tested. Predictions of knowledge of HIV status are nevertheless within 4% points of the ones obtained using the full data calibration, and have overlapping uncertainty intervals (Table 2).

**Figure 5.**
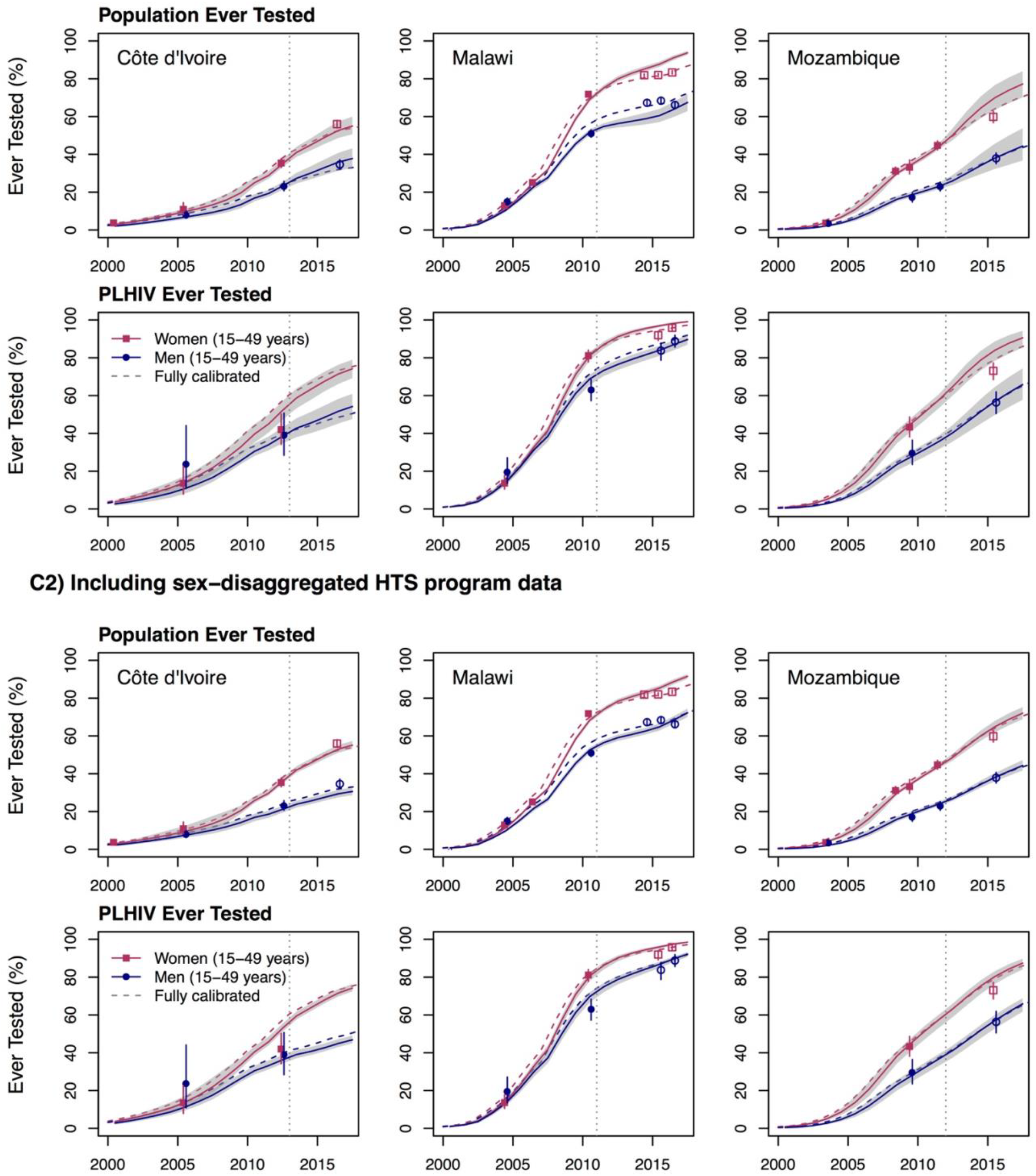
Out-of-sample predictions of F90 models calibrated to survey data from 2000-2012, including all available program data, for Côte d’Ivoire, Malawi, and Mozambique and model predictions. The top panel (C1) uses the overall (both sex combined) HIV Testing Services (HTS) program data whereas the bottom panel (C2) uses the sex-disaggregated HTS program data. Dashed lines represent predictions from the fully calibrated models. The vertical lines indicate the date of the last survey data estimates included in the fitting (to the right of the lines are the predictions). The shaded areas correspond to the 95% credible intervals of the posterior estimates. Points represent the survey estimates (Table 1) and empty symbols indicate that these outcomes were not included in the likelihood but are shown for cross-validation purposes. The vertical solid lines crossing these points correspond to the 95% confidence intervals of the survey estimates. (PLHIV: people living with HIV.)

On the other hand, the sex-disaggregated HTS program data generally increases the accuracy of the predictions –in both the overall population and among PLHIV (Figure 5, C2). In all countries, the F90 model’s predictions for the proportion of the population ever tested for HIV were ≤6% points different from the predictions obtained using the full data calibration (Table 2). Among PLHIV, all predictions had overlapping uncertainty intervals with those of the empirical survey estimates and differences were always less than 7% points. Predictions of the proportion of PLHIV aware of their status were also in very good agreement with those of the full data calibration, with differences of 3% points or less in all three countries.

## Discussion

Knowledge of HIV status is a key indicator to monitor progress, identify bottlenecks, and ultimately implement effective HIV responses. In this paper, we describe a new model that combines survey and HTS program data to estimate the “first 90” in SSA. We validated the F90 model through in-sample comparisons and our results demonstrate that it can accurately reproduce longitudinal sex-specific trends in HIV testing among the overall population and, more importantly, among PLHIV. Out-of-sample predictions of the fraction of individuals ever tested over a 4-to-6-year time horizon are also in good agreement with empirical survey estimates for PLHIV. When recent population-based surveys are not available, the accuracy of the F90 model predictions for the proportion of the population ever tested for HIV is improved by the addition of sex-disaggregated HTS program data. Importantly, our out-of-sample validations provided estimates of the fraction of PLHIV aware of their status that are consistent with the ones obtained using full data calibration and their uncertainty intervals overlap.

We compared our results to empirical estimates of awareness status among PLHIV. As expected, our F90 model predictions are higher than self-reported awareness status – even adjusted for presence of antiretroviral metabolites. For example, our model-based estimates of knowledge of HIV status are 58% (95%CrI: 53-61%; in 2017) in Côte d’Ivoire, 81% (95%CrI: 79-82%; in 2016) in Malawi, and 63% (95%CrI: 60-65%; in 2015), compared to 37% (not antiretroviral metabolites-adjusted), 76% (antiretroviral metabolites-adjusted), and 40% (antiretroviral metabolites-adjusted) of PLHIV who reported being aware of their HIV status in the 2017 Ivoirian, 2016 Malawian, and 2015 Mozambican surveys, respectively. The fundamental reason for the higher estimates of HIV status awareness is that the gap between the proportion of PLHIV ever tested, which are well reproduced by the model, and the proportion with knowledge of their status is constrained by the high rate of testing (i.e., any PLHIV who have ever been tested, but are not aware, must have been infected since the most recent HIV test). For the model’s predictions to be consistent with survey-based estimates of self-reported awareness, country-specific HIV incidence rates would need to be several-fold higher than the ones estimated by Spectrum/EPP and PHIA surveys and/or re-testing rates among PLHIV ever tested would have to be much lower than suggested by the survey and HTS program data. Other possible explanations include over-reporting of HIV testing history by survey respondents who were not aware, or very substantial levels of return of false-negative HIV test results, though such levels would have to be extremely high considering the high levels of re-testing.

The F90 model can be applied to countries with at least two population-based surveys that collected information on both HIV testing history and HIV prevalence. Because model predictions are expected to be more accurate over the short term, it is advisable to interpret with caution estimates produced for countries where the last population-based survey was conducted more than 5 years in the past. HTS program data on the number of tests performed should be carefully assessed to ensure that it accurately represents annual national testing volume among the population aged 15 years and over, and that it includes information from both private and public sectors. Sex-disaggregated HTS program data should be especially useful for countries without recent survey estimates of HIV testing histories. However, model predictions should be reasonably accurate, even if sex-disaggregated HTS data are unavailable, if the male/female testing ratios have remained relatively constant in recent years. Finally, it is advised to ensure some temporal degree of overlap between survey and program data to facilitate estimation of re-testing parameters. The latter is especially important if countries wish to examine additional model outputs of interest. For example, the model can provide information on the distribution of negative tests among first-time and repeat testers, and the distribution of positive tests among new diagnoses, retests among aware PLHIV, and retests among PLHIV on ART (Figure 6).

**Figure 6.**
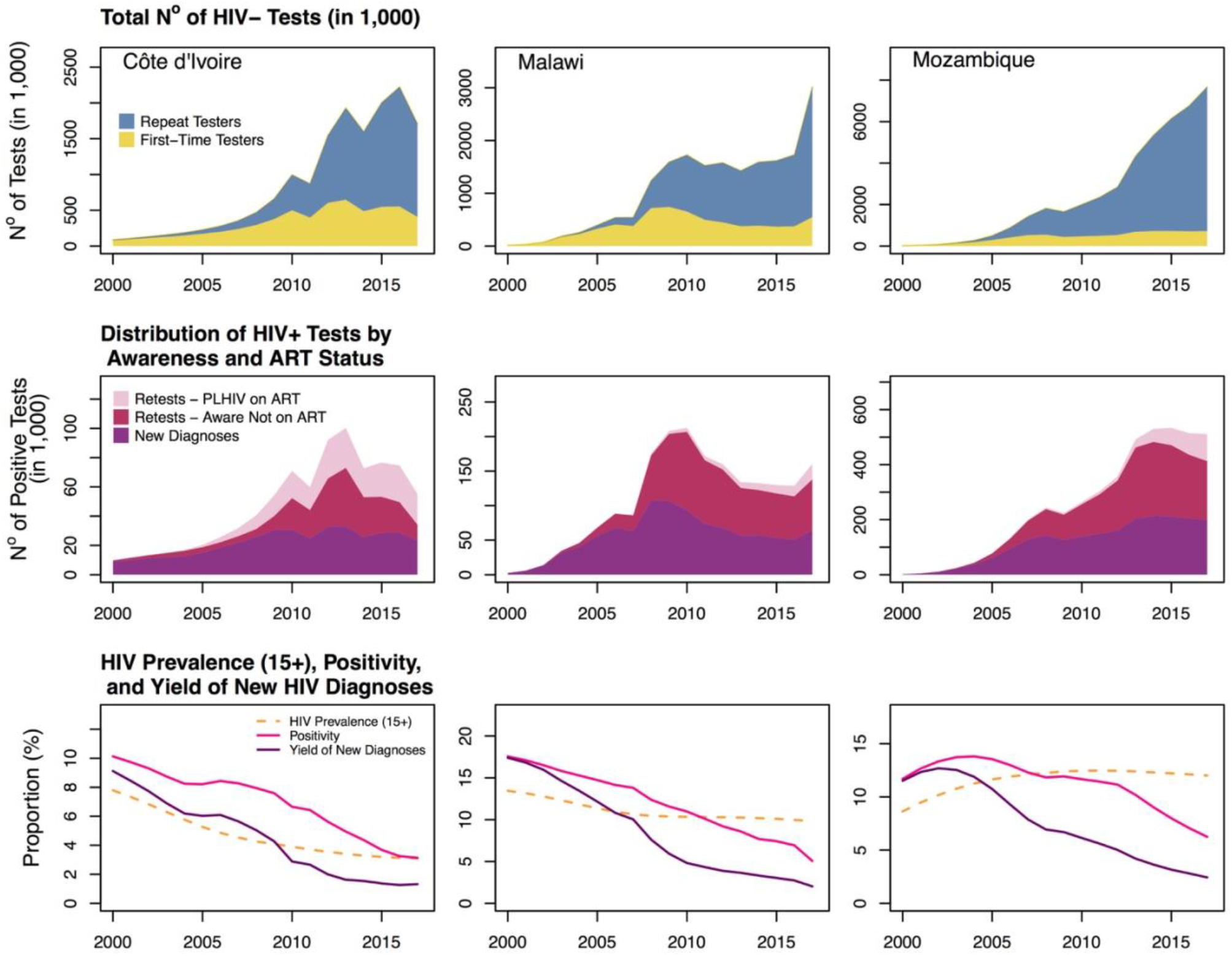
Model predictions of the distribution of the annual total number of HIV negative tests performed among first-time testers versus repeat-testers, distribution of HIV positive tests by awareness status and antiretroviral treatment (ART) status, and longitudinal trends in HIV testing positivity, yield of new diagnoses, and Spectrum/EPP’s estimates of HIV prevalence (aged 15+ years).

To facilitate model use, an online version was developed using the *RShiny* framework. Users can freely access the web-app (https://shiny.dide.imperial.ac.uk/shiny90/), review data sources, edit information, add new data, and run the model. It requires the users to provide a Spectrum/EPP projection file to use as input. Users can save their current analyses, perform sensitivity analyses, and export their results.

### Model limitations

Our proposed approach to estimate the “first 90” has several limitations, mainly due to data considerations. First, we assumed that self-reports of ever testing are accurate. Quantifying the sensitivity and specificity of those self-reports is difficult and their accuracy could differ by HIV status^[14]^. However, limited evidence suggest that testing histories are probably better reported than other potential indicators^[50]^ but incorrect reports of HIV testing history could result in underestimation of the “first 90”^[51]^. As evidence accrues on the sensitivity and specificity of those self-reports, adjustments for potential misclassification, if warranted, could be incorporated into the model. An additional source of uncertainty lies in the accuracy of HIV tests results provided back to HTS users. In the model, we assume that national HIV testing algorithms are accurate but some programs have reported suboptimal field sensitivity and specificity^[52–54]^. Second, published HTS program statistics usually relate to public sector programs and don’t necessarily reflect private sector testing, NGO testing programs, and self-testing. The latter poses additional challenges to the correct estimation of the number of HIV tests performed annually and difficulty in assessing trends over time in terms of positivity and yield of new diagnoses. We recommend sensitivity analyses to explore model robustness to assumptions regarding completeness of HTS program data. Thirdly, some national programs may have difficulties differentiating between tests performed on children aged less than 15 years from those in the modeled population. Data from Malawi suggest that children (<15 years) can comprise a small but non-negligible fraction (~16%) of overall testing volume^[55]^, though pediatric tests account for a substantially lower fraction of HIV-positive tests due to the low HIV prevalence in children, the results of effective prevention of mother-to-child transmission programs.

Regarding model structure and assumptions, a fourth limitation is that the current model implementation does not incorporate uncertainty in both the denominator of the “first 90” and the estimated ART coverage. This may result in an underestimation of uncertainty. Finally, the model does not currently disaggregate indicators by members of key populations (e.g., men who have sex with men, female sex workers, clients) or produce estimates of HIV diagnosis among children. Key populations are important to overall transmission dynamics in several countries^[56–59]^ and the sustainable control of HIV epidemics also hinges on also achieving the 90-90-90 targets in these groups^[60]^. The general framework outlined above could in theory be used to monitor awareness status for key populations, but additional challenges related to representativeness of key population surveys, among others, are expected^[13]^.

### Model strengths

Our proposed approach to estimate the proportion of PLHIV who know their status has several strengths. First, our model uses Spectrum outputs and is therefore fully consistent with other epidemiological data (e.g., sex and age-specific HIV incidence, prevalence, mortality) and programmatic outcomes (ART coverage). Second, it integrates routinely collected HTS program data with population-based surveys. This data triangulation enables monitoring of HTS’ effectiveness by providing estimates of annual new HIV diagnoses. Third, our approach attempts to overcome the limitations of self-reported knowledge of HIV status by pooling information on ART coverage, HIV re-testing rates, and HIV incidence to estimate how many PLHIV acquired their infection after their last HIV-negative test. Finally, the current framework enables us to further refine the F90 model and its assumptions as more granular program data become available (e.g., age-stratified HIV testing program data) and provides a foundational framework for future work to incorporate data about HIV testing and diagnosis into estimates of HIV incidence trends.

## Conclusions

Identifying the proportion of PLHIV who know their status is challenging and relying solely on self-reported estimates of HIV awareness could be misleading. The aim of our F90 model is to triangulate different data sources to improve the accuracy of the “first 90” indicator. Beyond the estimation of HIV status knowledge, the F90 model also produces estimates of annual number of new HIV diagnoses. Such information can help countries improve the effectiveness of their HIV testing programs and assist them in reaching the first 90 target by 2020.

## AUTHORS’ CONTRIBUTION

CH, DB, JWE, KM, MMG, and RB conceived and designed the model. AG, AJ, AK, CD, CLD, FM, JWE, KM, and MMG obtained, administered, and processed the different databases. AG, AJ, CD, JE, JL, KM, MCB, ME, and MMG contributed to model development and/or revisions. AG, CD, CLD, JWE, and MMG performed the analyses and all authors contributed to results interpretation. MMG drafted the manuscript and all authors critically reviewed it for important intellectual content. All authors approved the final version.

## ACKNOWLEDGMENTS

We are grateful to Shelley Clark for her comments and suggestions and to Alexandra Hill and Martin Eden for help in developing the Shinny 90 app. Also to thank are Roma Bhatkoti, Drew Voetsch, Joshua Salomon, John Stover, Ray Shiraishi and other participants of the *UNAIDS Reference Group on Modelling, Estimates, and Projections* for useful feedback on an earlier version of the proposed modeling framework.

We acknowledge funding from the *Steinberg Fund for Interdisciplinary Global Health Research* (McGill University) and the *Bill and Melinda Gates Foundation*. MMG’s research program is funded by a career award from the *Fonds de recherche du Québec – Santé*. JWE was supported by UNAIDS and the *Bill and Melinda Gates Foundation*.

