## Supplemental Materials for "National HIV testing and diagnosis coverage in sub-Saharan Africa: a new modeling tool for estimating the “first 90” from program and survey data"

**Text S1**. Full description of the model's parameterization for HIV testing rates and elicited prior distributions.

*General parameterization*

The rate at which individuals are tested for HIV varies by calendar time ($t$), sex ($k$), age ($i$), HIV testing history and awareness status ($u$), and for PLHIV, CD4+ cell count category ($s$). As stated in the main text, it takes the following form:

$$\tau_{kius}(t) = b(t)M_k(t)A_{ki}F_u(t) + \rho_k(t)OI_s$$

*Testing rates for referent group (b(t))*

The *b(t)* rate is parameterized such that the referent group is that of women aged 15-24 years. It is assumed that *b(t)* is equal to zero in 1995 and then increases linearly to $\beta_{2000}$ in the year 2000. After that time, testing rates are assumed to follow a first order random walk process (RW1) with annual time steps. The complete specification is:

$$b(t) = \begin{cases} 0;\ if\ t \leq 1995 \\ \dfrac{\beta_{2000}}{5} \times (t - 1995);if\ 2000 < t > 1995 \\ \beta_{2000};if\ t = 2000 \\ log\big(b(t)\big) \sim N\big(log\big(b(t-1)\big), \sigma_{rate}\big);if\ t > 2000 \end{cases}$$

This specification is completed with the following prior distributions (below). For $\beta_{2000}$, we assumed that testing rates would be 0.01 per year, with 95% of the prior distribution's density between the rate of 0 and 0.02 per year. For the standard deviation of the RW1 process, we chose to fix it such that, if the baseline testing rate at time *t* was 0.25 per year (once per 4 years on average), 95% of the distribution's density for the next year's rate would be within plus or minus 50% of that value.

$$log(\beta_{2000}) \sim N(log(0.001), 0.25)$$

$$\sigma_{rate} = 0.35$$

It should be mentioned that the BFGS calibration algorithm we chose solves unconstrained optimization problems (i.e., it takes input parameter values between –∞ and ∞). To restrict parameter values o epidemiologically appropriate scales, logarithmic (as above) and logit transformations are used, when appropriate.

*Testing rates by sex ($M_k(t)$)*

As the referent rate is defined for females aged 15-24 years, the term $M_k(t)$ is introduced to allow a different testing rate for males aged 15-24 years. Further, with the scale-up of prevention of mother-to-child transmission program in sub-Saharan Africa, we expect this ratio to change with time between 2005 and 2012, as detailed below.

$$M_k(t) = \begin{cases} \omega_{2005};if\ k = 1\ and\ t \leq 2005 \\ \omega_{2005} + \dfrac{(\omega_{2012} - \omega_{2005})}{7} \times (t - 2005);if\ k = 1\ and\ 2005 > t < 2012 \\ log(\omega_{2012}) \sim N(log(\omega_{2005}), \sigma_{rr});if\ k = 1\ and\ t \geq 2012 \\ 1;if\ k = 2 \end{cases}$$

where $\omega_{2005}$ and $\omega_{2012}$ are the rate ratios for male testing in 2005 and 2012, respectively. To define appropriate prior distributions for these parameters, we first estimated the male-to-female ratios for reporting having been tested for HIV in the last 12 months (and received the results). Based on all available nationally-representative population-based surveys conducted in sub-Saharan Africa (total of 110 surveys; N = 375,302), we calculated using a random effect meta-analysis this pooled rate-ratio, estimated at 0.6. This value was chosen as the median of the prior for the 2005 male testing ratio ($\omega_{2005}$). Since almost all survey estimates were smaller than one, we bounded the ratio between 0 and 1.10 using a logit transformation. The prior for this parameter thus correspond a distribution where 95% of its density is between the ratios of 0.07 and 1.05. Finally, the standard deviation of the RW1 process of the 2012 rate ratio ($\omega_{2012}$) was chosen such that if $\omega_{2005}$ has its prior value of 0.6, $\omega_{2012}$ will have 95% of its distribution in the 0.36-1.00 interval.

$$\omega_{2000} \sim logit^{-1}\left(N(logit\left(\frac{0.6}{1.1}\right), 1.46)\right) \times 1.1$$

$$\sigma_{rr} = 0.25$$

*Testing rates by age ($A_{ki}$)*

We expect that testing rates, not only differ by sex, but also change as a function of age ($i$). To take this relationship between age and testing rates into consideration, we specified time-invariant sex-specific rate ratios ($A_{ki}$) for four different age group: 15-24 years old ($i=1$), 24-34 years ($i=2$), 35-49 years ($i=3$), and 50+ ($i=4$). Few surveys collected information on men and women aged 50+ years. It is therefore not possible to estimate a rate ratio for this age group for most countries. To account for potential changes in testing ratios among the 50+ years old, we conduct meta-analyses of 124 and 81 surveys for females (N = 156,566) and males (N = 101,557), respectively. For females, we estimated a pooled random-effect rate ratios of testing in the last 12 months between the 40-44 and 45-49 age group as the overwhelming majority of surveys did not collect information on women aged 50+ years. For males, we compared testing among the 35-49 and 50-59 age groups. For both sexes, the meta-analyzed rate ratio was 0.81. This was used to parameterize $A_{ki}$ the following way:

$$A_{ki} = \begin{cases} 1.00; if\ i = 1 \\ \alpha_{ki}; if\ i \in \{2,3\} \\ 0.81 \times \alpha_{k=1,i=3}; if\ \ i = 4 \\ 0.81 \times \alpha_{k=2,i=3}; if\ \ i = 4 \end{cases}$$

To help with model calibration, we restricted the parameter space by assuming that all age-specific rate ratios would be bound between 0.10 and 6.00. Specifically, the prior distributions for testing ratios by age were given a mean of 1.00 and 95% of their distribution's density would lie in the 0.15 to 5.00 interval, as described below.

$$\alpha_{k,i\in\{2,3\}} \sim logit^{-1}\left(N(logit\left(\frac{1-0.1}{6-0.1}\right), 1.685\right) \times (6.00 - 0.1) + \ 0.1$$

*Testing rates by HIV testing history and HIV status ($F_u(t)$)*

Potential differences in testing rates as a function of HIV status and past testing history are considered using the $F_u(t)$ parameter. Here, the '$u$' subscript indicated if an individual is: HIV-susceptible never tested ($u$=1), HIV-susceptible previously tested ($u$=2), PLHIV never tested ($u$=3), PLHIV unaware ever tested ($u$=4), PLHIV aware not on treatment ($u$=5), and PLHIV on ART ($u$=6). This takes the following general form:

$$F_u(t) = \begin{cases} 1.00; if\ u = 1 \\ RR_{test}(t); if\ u = 2 \\ RR_{unaware}; if\ u = 3 \\ RR_{test}(t) \times RR_{unaware}; if\ u = 4 \\ RR_{aware}(t); if\ u = 5 \\ RR_{art}; if\ u = 6 \end{cases}$$

To allow for changes in re-testing rates among HIV-susceptible individuals who have been previously tested ($u$=2), we allowed the rate ratio ($RR_{test}$) to vary between 2010 and 2015, with piecewise linear interpolation in between these years. (There are few HTS program data available prior to 2010 and we held retesting ratios constant before that date.) The rate ratio of re-testing among HIV-susceptible individuals is equal to:

$$RR_{test}(t) = \begin{cases} \rho_{2010}; if\ t \leq 2010 \\ \rho_{2010} + \frac{(\rho_{2015} - \rho_{2010})}{5} \times (t - 2010); if\ 2010 > t < 2015 \\ log(\rho_{2015}) \sim N(log(\rho_{2010}), \sigma_{test}); if\ t \geq 2015 \end{cases}$$

Similarly, rate ratios of re-testing among those aware of their status and not on treatment ($u$=5) were also allowed to vary using RW1 process with annual time steps, taking the following form:

$$RR_{aware}(t) = \begin{cases} \delta_{2010}; if\ t \leq 2010 \\ log\big(RR_{aware}(t)\big) \sim N\big(log\big(RR_{aware}(t-1)\big), \sigma_{aware}\big); if\ t > 2010 \end{cases}$$

The prior distributions for the parameters in $F_u(t)$ are explained below. For the prior retesting rate ratio ($\rho_{2010}$) among HIV-susceptible with a history of HIV testing ($u$=2), we performed a meta-analysis of three studies described in Table S1 to inform the mean of the parameter's distribution. Due to potentially high heterogeneity between countries, this parameter was given a rather uninformative distribution. Specifically, we bound the distribution between 0.95 and 8.00, with 95% of the distribution's density between the values of 1.08 and 5.00. The rate ratio for this group in 2015 ($\rho_{2015}$) is modeled as a RW1 process with a standard deviation of 0.25. This entails that, for a previous rate ratio of 1.00, 95% of the prior's density for the rate ratio would be within the 0.6 to 1.6 interval. The prior distribution for the rate ratio for testing among PLHIV unaware ($RR_{unaware}$) was given a mean of 1.00 and was bounded between the ratios of 0.05 and 1.95. Within these bounds, 95% of its distribution's density would be between the ratios of 0.10 and 1.90. For the testing ratio among PLHIV aware of their status not on treatment ($u$=5), we assumed that the mean ratio would be 1.50 and bounded between rate ratios of 0.00 and 8.00, with 95% of its distribution's density between the ratio of 0.14 and 6.00. Finally, the prior distribution for the rate ratio for individuals on ART ($u$=6) was given a prior that assumed that the ratio would be bound between 0 and 1 (see Table S2), with a mean of 0.25 and 95% of its distribution's density between the value of 0.01 and 0.90. These distributions are descried below.

$$\rho_{2010} \sim logit^{-1}\left(N(logit\left(\frac{1.93 - 0.95}{8 - 0.95}\right), 1.084\right) \times (8 - 0.95) + \ 0.95$$

$$\sigma_{test} = 0.25$$

$$RR_{unaware} \sim logit^{-1}(N(logit(0.5), 1.1) \times (1.95 - 0.05) + \ 0.05$$

$$\delta_{2010} \sim logit^{-1}\left(N(logit\left(\frac{1.5}{8}\right), 1.31\right) \times 8$$

$$\sigma_{aware} = 0.25$$

$$RR_{art} \sim logit^{-1}(N(logit(0.25), 1.68)$$

*Testing from symptomatic HIV infection ($\rho_k(t)OI_s$)*

The specification of the HIV testing rate $\tau_{kius}(t)$ has so far not considered testing that may occur, among those PLHIV not on treatment, because of HIV/AIDS-related symptoms. For individuals experiencing such symptoms, we consider the additive rate defined by $\rho_k(t)OI_s$. The term $OI_s$ is the time-invariant incidence of opportunistic infections for PLHIV in CD4+ cell count category $s$ [1, 2]. Progression through those different CD4+ cell count categories are tracked by Spectrum. We assume that the sex-specific proportion of opportunistic infections tested for HIV varied in time, proportionally to that of the sex-specific ART coverage, and can reach a maximum of 95%, as specified below.

$$\rho_k(t) = max\{ART_k(t) \times \kappa; 0.95\}$$

$$OI_s = \begin{cases} 0; if\ s = 0\ (CD4 \geq 350\ cells\ \mu L^{-1}) \\ 0.27;\ if\ s = 1\ (CD4\ 200 - 349\ cells\ \mu L^{-1}) \\ 0.90;\ if\ s = 2\ (CD4\ < 200\ cells\ \mu L^{-1}) \end{cases}$$

The parameter $\kappa$ ensures that the proportion of opportunistic infections is scaled-up proportionally to sex-specific ART coverage, as obtained from Spectrum/EPP. The prior distribution for $\kappa$ has a mean of 1.00 and is bounded by 0.25 and 1.75, with 95% of the prior distribution's density between the values of 0.30 and 1.75.

$$\kappa \sim logit^{-1}(N(logit(0.5), 1.75) \times (1.75 - 0.25) + 0.25$$

### F90 model equations

The dynamic system formed by the different compartments of the F90 model can be represented by the set of six equations below. All individuals enter the population at 15 years of age, informed by Spectrum/EPP demographic parameter, and are assumed to have never been tested $E_{k,i=1}(t)$ if susceptible or living with HIV but untreated $L_{k,i=1}(t)$, unless they are already living with HIV and on ART $H_{k,i=1}(t)$ in which case they go to the corresponding compartment (i.e., u=6). Aging is also informed by Spectrum through the $Aged_{kisu}(t)$ parameter. Individuals are then subjected to age- and sex-specific HIV incidence $\lambda_{ki}(t)$, natural mortality $\upsilon_{ki}(t)$, and HIV testing $\tau_{kis,u}(t)$. Once HIV infected, individuals follow the disease's natural progression (tracked by Spectrum/EPP) and can die of HIV-related causes at rate $\epsilon_{ksu}$. As stated in the main text, the F90 model is concerned with estimating $\tau_{kis,u=1}(t)$ and all other parameters are taken directly from Spectrum/EPP (i.e., $E_{ki}(t)$, $L_{ki}(t)$, $H_{ki}(t)$, $Aged_{kisu}(t)$, $\lambda_{ki}(t)$, $\upsilon_{ki}(t)$, $\epsilon_{ksu}$), including progression through the different CD4+ cell count categories (not shown in the equations below).

*HIV-susceptible never tested (u=1)*

$$\frac{dX_{kis,u=1}(t)}{dt} = E_{ki}(t) + Aged_{kis,u=1}(t) - \left(\tau_{kis,u=1}(t) + \lambda_{ki}(t) + \upsilon_{ki}(t)\right)X_{kis,u=1}(t)$$

*HIV-susceptible ever tested (u=2)*

$$\frac{dX_{kis,u=2}(t)}{dt} = Aged_{kis,u=2}(t) + \tau_{kis,u=1}(t)X_{kis,u=1}(t) - (\lambda_{ki}(t) \ + \upsilon_{ki}(t))X_{kis,u=2}(t)$$

*PLHIV never tested (u=3)*

$$\frac{dX_{kis,u=3}(t)}{dt} = \ L_{ki}(t) + Aged_{kis,u=3}(t) + \ \lambda_{ki}(t)X_{kis,u=1}(t) - (\tau_{kis,u=3}(t) + \upsilon_{ki}(t) + \epsilon_{ksu})X_{kis,u=3}(t) - c_{kis,u=3}(t)$$

*PLHIV ever tested and unaware (u=4)*

$$\frac{dX_{kis,u=4}(t)}{dt} = Aged_{kis,u=4}(t) + \lambda_{ki}(t)X_{kis,u=2}(t) - (\tau_{kis,u=4}(t) + \upsilon_{ki}(t) + \epsilon_{ksu})X_{kis,u=4}(t) - c_{kis,u=4}(t)$$

*PLHIV aware but not on treatment (u=5)*

$$\frac{dX_{kis,u=5}(t)}{dt} = Aged_{kis,u=5}(t) + \tau_{kis.u=3}(t)X_{kis,u=3}(t) + \tau_{kis,u=4}(t)X_{kis,u=4}(t) + \gamma_{ks}(t)X_{kis,u=6}(t) - (\eta_{ks}(t) + \upsilon_{ki}(t) + \epsilon_{ksu})X_{kis,u=5}(t)$$

*PLHIV on treatment (u=6)*

$$\frac{dX_{kis,u=6}(t)}{dt} = H_k(t) + Aged_{kis,u=6}(t) + c_{kis,u=3}(t) + c_{kis,u=4}(t) + \eta_{ks}(t)X_{kis,u=5}(t) - (\gamma_{ks}(t) + \upsilon_{ki}(t) + \epsilon_{ksu})X_{kis,u=6}(t)$$

In rare instances, rapid ART scale-up could deplete the compartment of PLHIV aware not on treatment (*u*=5). If this is the case, individuals newly initiated on ART are taken from the unware compartments ($u \in \{3,4\}$ ), in a proportion relative to the size of PLHIV ever/never tested, and are assumed to be immediately put on ART after diagnosis. The following constraint is used to define the parameter $c_{kis,u\in\{3,4\}}(t)$ that describes the number of immediately ART-initiated individuals at time *t*.

$$Constraint_{kis}(t) = \ \left(\eta_{ks}(t)X_{kis,u=5}(t)\right) dt - (Cov_{kis}(t+dt) - Cov_{kis}(t))$$

where $Cov_{kis}(t)$ is the Spectrum/EPP estimates of the number of PLHIV on ART at time *t*.

$$c_{kis,u=3}(t) = \begin{cases} 0; if\ Constraint_{kis}(t) \geq 0 \\ \begin{pmatrix} X_{kis,u=6}(t+dt) - X_{kis,u=6}(t)) - \\ \left(\eta_{ks}(t)X_{kis,u=5}(t)\right) \end{pmatrix} \times \frac{X_{kis,u=3}(t)}{\sum_{u\in\{3,4\}} X_{kisu}(t)}; if\ Constraint_{kis}(t) < 0 \end{cases}$$

$$c_{kis,u=4}(t) = \begin{cases} 0; if\ Constraint_{kis}(t) \geq 0 \\ \begin{pmatrix} X_{kis,u=6}(t+dt) - X_{kis,u=6}(t)) - \\ \left(\eta_{ks}(t)X_{kis,u=5}(t)\right) \end{pmatrix} \times \frac{X_{kis,u=4}(t)}{\sum_{u\in\{3,4\}} X_{kisu}(t)}; if\ Constraint_{kis}(t) < 0 \end{cases}$$

**Text S2**. Definition of the F90 model likelihood function and other model outputs.

Depending on data availability, the model can be fitted to: 1) the proportion of individuals self-reporting having ever tested for HIV for calendar year *t*, stratified by age, sex, and HIV status (if available); 2) the annual total number of HIV tests performed for calendar year *t* (stratified by sex or not); and 3) the annual total number of HIV tests found to be positive for calendar year *t* (stratified by sex or not). Because of the model's relatively simple structure and tractability of its likelihood formulas, a Bayesian calibration framework was chosen for its computational efficiency. Further, the motivation for a likelihood-based approach for inference is to account for the sampling variability in the data and weight data sources relative to their precision.

*Proportion ever tested*

The likelihood of the model parameters $\theta$ given the survey data about the proportion ever tested ($S_{kist}$), and available for calendar years contained in the vector $T^a$, can be written as follows.

$$\mathcal{L}(\theta|S_{kis,t\in\{Y^a\}}) = \prod_{t\in\{Y^a\}}\prod_k\prod_i\prod_s \binom{n_{tkis}}{x_{tkis}} \zeta_{tkis}^{x_{tkis}}(1-\zeta_{tkis})^{n_{tkis}-x_{tkis}}$$

where $\zeta_{tkis}$ is the model-predicted proportion of the population having ever been tested for HIV, $x_{tkis}$ is the survey-weighted number of individuals that report having ever been tested for HIV out of an effective sample size of $n_{tkis}$ (adjusted for survey design effects) in a survey conducted at time *t*, of sex *k*, of age *i*, and HIV status *s*. If the survey did not include HIV serology, then the proportion having ever been tested for HIV are predicted for all persons, both HIV positive and HIV negative. Because all the survey types (e.g., AIS, DHS, MICS, PHIA, etc.) used for model calibration are nationally representative population-based surveys with similar design and questionnaire, we only consider the above specification of the likelihood function.

The model predicted proportion of the population ever tested for HIV is calculated as follow, where $X_{kisu}(t)$ is the number of individuals of sex (*k*), age (*i*), HIV testing history and awareness status (*u*), and, for PLHIV, CD4+ cell count (*s*).

$$\zeta_{kis}(t) = \frac{\sum_k\sum_i\sum_s X_{kis,u\in\{2,4,5,6\}}(t)}{\sum_k\sum_i\sum_s\sum_u X_{kisu}(t)}$$

*Programmatic data on number of HIV tests*

The likelihoods of the model parameters $\theta$ given the data on the number of HIV tested for ($T_{kt}$) and number of positive tests ($P_{kt}$) for sex *k* performed at time *t* (and available for the years in vectors $Y^b$ and $Y^c$, respectively) is the product of two functions:

$$\mathcal{L}\left(\theta\middle|T_{k,t\in\{Y^b\}}\right) = \prod_{t\in\{Y^b\}}\prod_k \frac{1}{\sqrt{2\pi(T_{kt}*0.05)^2}} e^{\frac{-(T_{kt}-\mu_k(t))}{2(T_{kt}*0.05)^2}}$$

$$\mathcal{L}(\theta|P_{k,t\in\{Y^c\}}) = \prod_{t\in\{Y^c\}}\prod_k \frac{1}{\sqrt{2\pi(P_{kt}*0.10)^2}} e^{\frac{-(P_{kt}-\varphi_k(t))}{2(P_{kt}*0.10)^2}}$$

where $T_{tk}$ is the total number of tests performed, and reported for calendar years in $Y^b$, for sex *k*; $\mu_k(t)$ is the model-predicted number of tests performed; $P_{kt}$ is the total number of HIV positive tests in year *t* and reported for calendar years in $Y^c$; $\varphi_k(t)$ is the model-predicted number of HIV positive tests. Because there is no sampling variance associated with HTS program data, we assumed that the standard errors for these outcomes would be equal to 5% and 10% of the total for the number of positive tests and overall number of tests, respectively. This ensures that the model can reproduce the temporal patterns in the HTS program data but also allows some trade-off between fitting the survey and HTS program data if the two data sources are not fully consistent.

The total model-predicted number of HIV tests performed during year *t* can be estimated using:

$$\mu(t) = \int_{t-1}^{t} \sum_{k} \sum_{i} \sum_{s} \sum_{u} \tau_{kius}(t) X_{kius}(t) dt$$

And the total model-predicted number of tests found to be positive during year *t* as:

$$\varphi(t) = \int_{t-1}^{t} \sum_{k} \sum_{i} \sum_{s} \sum_{u \in \{3,4,5,6\}} \tau_{kius}(t) X_{kius}(t) dt$$

The overall model log-likelihood is obtained by summing the log-likelihood of the three likelihood functions. The relative weight of each data sources used in model calibration is thus a function of the number of data points (and effective sample size of the survey data). If there is no HTS program data to calibrate the model to, only the survey data on the proportion of the population reporting having ever been tested is used.

*Other model outputs*

Estimates of the proportion of PLHIV with knowledge of status *A(t)* can be calculated as follow.

$$A_{ki}(t) = \frac{\sum_k \sum_i \sum_s X_{kis,u \in \{5,6\}}(t)}{\sum_k \sum_i \sum_s \sum_u X_{kis,u \in \{3,4,5,6\}}(t)}$$

**Text S3**. Comparisons of annual number of HIV tests performed with self-reports of HIV testing in the last year.

Population-based surveys which include a HIV testing history module often query if individuals have ever been tested for HIV and if their last test occurred in the past year. Responses from this latter survey item could overestimate the proportion of individuals tested in the past year due to telescoping bias[3-5]. This bias could occur if survey respondents lengthen the recall period beyond the last 12 months. To assess the magnitude of this potential telescoping bias, we compare estimates of the number of HIV tests calculated from survey estimates to programmatic data on the reported number of HIV tests performed in the year before the survey (or closest available date). The survey-based number of HIV tests was obtained by converting the proportion reporting an HIV test in the past year to a rate and multiplying this rate by the appropriate population denominator (population aged ≥15 years). The results show that for almost all countries with suitable data, the survey-based extrapolations of the number of HIV tests performed overestimate the ones obtained from HIV testing programs (Figures S1 and S2). It is unlikely that this overestimation could be explained by the shortcomings of this analysis, which include: 1) small mismatch between the calendar year for which programs report number of tests and survey recall period, b) imperfect concordance between survey population of individuals aged 15-49 or 15-59 and population aged ≥15 years being tested, and c) potential underreporting of HIV tests performed by private sector.

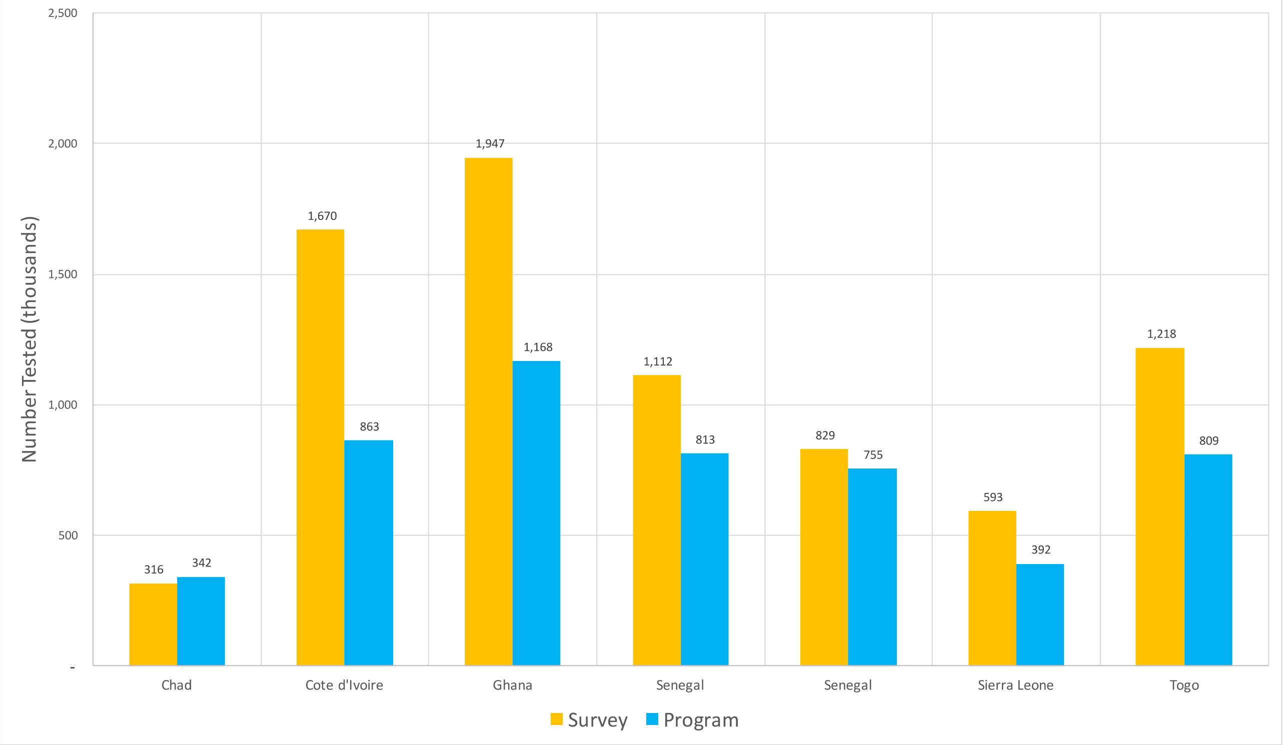


**Figure S1.** Comparisons of survey-based extrapolations of the number of HIV tests performed and the number of HIV tests reported by countries in West and Central Africa.

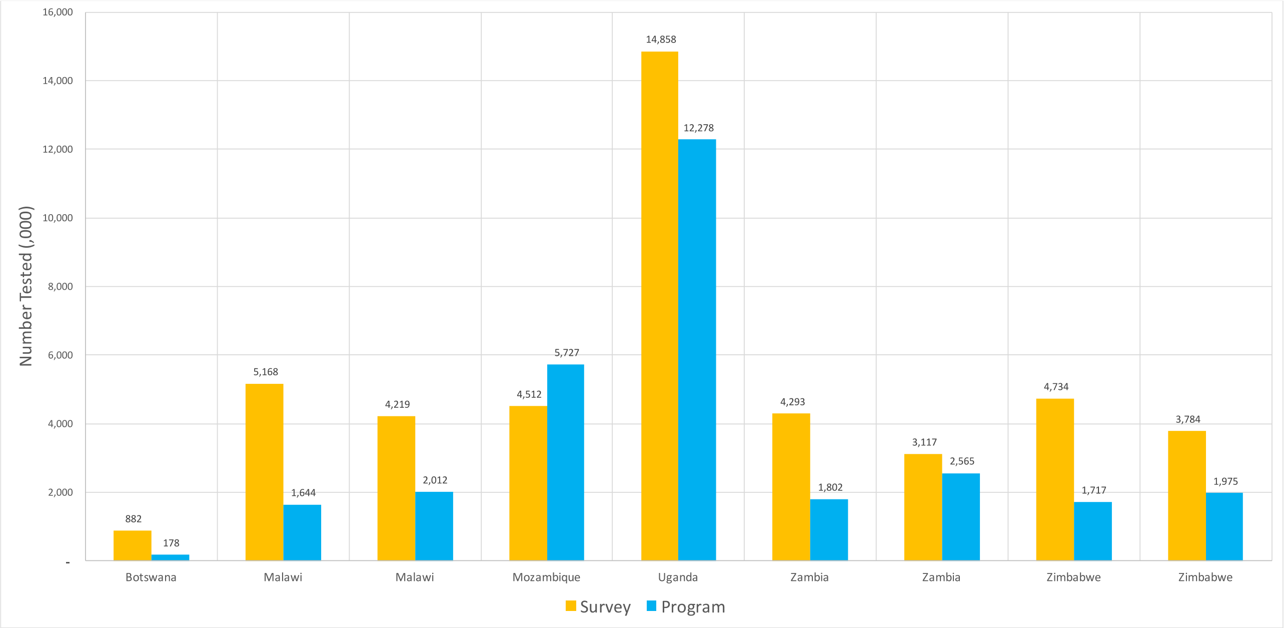


**Figure S2.** Comparisons of survey-based extrapolations of the number of HIV tests performed and the number of HIV tests reported by countries in East and South Africa.

**Table S1.** Summary of findings from main studies comparing testing rates relative to testing history and awareness status.

| Author, publication year | Study period | Country | Design | Sample size | Population | Testing services offered / available | Exposure(s) | Outcome(s) | Effect size measure |
|---|---|---|---|---|---|---|---|---|---|
| Floyd, 2013[6] | 2007-08, 2008-09, and 2009-10 | Malawi | ♦Cross-sectional design ♦Serological surveys (3 waves) | 41,353 | Rural population, ≥15 years old | HIV testing | Result of the previous HIV test (positive vs. negative) | Accepting HIV testing | RR of accepting HIV testing (previously tested positive vs. previously tested negative): **0.69** (95% CI: 0.66-0.72) |
| Ng'ang'a, 2014[7] | 2012-13 | Kenya | ♦Cross-sectional design ♦Population-based household survey | 13,655 | 15-64 years old | HBTC | History of HIV testing (ever testers vs. never testers) | Accepting HBTC* | RR of accepting HBTC (ever testers vs. never testers): **0.94** (95% CI: 0.89-0.99) |
| Dalal, 2011[8] | 2007 | South Africa | ♦Cross-sectional design ♦Pre-intervention/post intervention study | 912 | Patients from the general outpatient clinic, 18-49 years old | PITC | History of HIV testing (ever testers vs. never testers) | HIV test uptake | RR of using testing and counselling services (ever testers vs. never testers): **1.37** (95% CI: 1.03-1.88) |
| South, 2013[9] | 2006-07 | Tanzania | ♦Cross-sectional design ♦Survey | 9,641 | Rural population, ≥15 years old | VCT | History of VCT use (ever users vs. never users) | VCT uptake | RR of using VCT (ever testers vs. never testers): **2.16** (95% CI: 1.96-2.38) |
| Reniers, 2009[10] | 2005-2006 | Zimbabwe | ♦Cross-sectional design ♦Serological survey (DHS) | *Not provided* | *Not provided* | HIV testing | History of HIV testing (ever testers vs. never testers) | Accepting to test for HIV* | RR of accepting to test for HIV (ever testers vs. never testers): **0.98** |
| Reniers, 2009[10] | 2005 | Senegal | ♦Cross-sectional design ♦Serological survey (DHS) | *Not provided* | *Not provided* | HIV testing | History of HIV testing (ever testers vs. never testers) | Accepting to test for HIV* | RR of accepting to test for HIV (ever testers vs. never testers): **0.94** |

| | | | | | | | | | |
|---|---|---|---|---|---|---|---|---|---|
| Reniers, 2009[10] | 2004 | Cameroon | ♦Cross-sectional design ♦Serological survey (DHS) | *Not provided* | *Not provided* | HIV testing | History of HIV testing (ever testers vs. never testers) | Accepting to test for HIV* | RR of accepting to test for HIV (ever testers vs. never testers): **0.95** |
| Reniers, 2009[10] | 2004 | Malawi | ♦Cross-sectional design ♦Serological survey (DHS) | *Not provided* | *Not provided* | HIV testing | History of HIV testing (ever testers vs. never testers) | Accepting to test for HIV* | RR of accepting to test for HIV (ever testers vs. never testers): **1.01** |
| Reniers, 2009[10] | 2004 | Lesotho | ♦Cross-sectional design ♦Serological survey (DHS) | *Not provided* | *Not provided* | HIV testing | History of HIV testing (ever testers vs. never testers) | Accepting to test for HIV* | RR of accepting to test for HIV (ever testers vs. never testers): **0.95** |
| Reniers, 2009[10] | 2003 | Ghana | ♦Cross-sectional design ♦Serological survey (DHS) | *Not provided* | *Not provided* | HIV testing | History of HIV testing (ever testers vs. never testers) | Accepting to test for HIV* | RR of accepting to test for HIV (ever testers vs. never testers): **0.97** |
| Isingo, 2012[11] | 2003-04 and 2006-07 | Tanzania | ♦Longitudinal design ♦Linked serological surveys (2 waves) | 18,417 | Rural population, ≥15 years old | VCT | History of VCT use (ever users vs. never users) | Accepting VCT | RR of using VCT (ever testers vs. never testers): **2.20** (95% CI: 2.01-2.43) |

95%CI: 95% confidence intervals; DHS: demographic and health survey; HBTC: home-based testing and counselling; PITC: provider-initiated testing and counselling; RR: relative risk; VCT: voluntary counselling and testing.
* These studies performed HIV tests for research purposes without disclosing the results to the participants. The incentives for accepting HIV testing in such instances could be different than if HIV testing was performed for clinical purposes.

**Table S2**. Estimating the risk ratio of re-testing within the past year among people living with HIV (PLHIV) aware on antiretroviral therapy (ART) compared to PLHIV aware but not on ART using data from the 2010 Malawi Demographic and Health Survey (DHS).

| Sex | Risk ratio | 95% CI* |
|---|---|---|
| Women | 0.65 | (0.47, 0.92) |
| Men | 0.55 | (0.33, 0.94) |
| *Overall* | 0.61 | (0.45, 0.81) |

95%CI: 95% confidence intervals.
Our analyses were restricted to respondents with a positive HIV tests (biomarker) in the survey. We then compared the proportion of respondents that reported having been tested for HIV in the last 12 months between those that reported being currently on antiretroviral therapy (ART) to the proportion of those tested in the last 12 months that were aware (self-reported) of their HIV-positive status but not on ART. We calculated this estimate overall, as well as stratified by sex. The 95%CI are obtained through 1,000 bootstrap replicates, where the resampling unit is the cluster.
Note that the estimated risk ratio above does not equal to the model's rate ratio for those on ART (i.e., $F_{u=6}(t)$) since the latter does not take into account HIV testing resulting from incident opportunistic infections.

**Table S3**. Comparisons of the posterior distributions of different model estimates obtained using the *Laplace* approximation, *Sampling Importance Resampling* (SIR), and *Incremental Mixture Importance Sampling* (IMIS) for Côte d'Ivoire, Malawi, and Mozambique.

| Outcomes and Methods | Côte d'Ivoire (95%CrI) | Malawi (95%CrI) | Mozambique (95%CrI) |
|---|---|---|---|
| **Ever tested (%) in 2017** | | | |
| Laplace approximation | 42.6% (41.9-43.7%) | 78.3% (77.1-79.6%) | 56.0% (54.8-57.4%) |
| SIR* | 42.6% (41.7-43.5%) | 78.3% (77.1-79.5%) | 55.8% (54.7-57.3%) |
| IMIS | 42.6% (41.9-43.4%) | 78.3% (77.3-79.3%) | 56.0% (54.8-57.2%) |
| **PLHIV aware (%) in 2017** | | | |
| Laplace approximation | 57.8% (52.5-61.3%) | 84.4% (82.9-85.8%) | 71.5% (68.8-73.3%) |
| SIR* | 58.2% (53.4-62.1%) | 84.3% (83.0-85.7%) | 71.1% (69.0-73.4%) |
| IMIS | 57.9% (53.8-60.7%) | 84.4% (83.1-85.5%) | 71.3% (69.3-73.0%) |

95%CrI: 95% credible interval.
SIR: sampling importance resampling; IMIS: Incremental Mixture Importance Sampling.
*The proposal distribution for SIR is based on a multivariate *t* distribution with 2 degrees of freedom, itself approximated using Laplace.

**Table S4**. List of main F90 model outputs and available disaggregation levels.

| Outputs | Available disaggregation levels |
|---|---|
| **Programmatic** | |
| Number of HIV tests. | Age, sex, calendar year. |
| Number of negative tests. | Age, sex, testing history, calendar year. |
| Number of positive tests. | Age, sex, testing history, calendar year. |
| Number of new diagnoses. | Age, sex, testing history, calendar year. |
| Number of repeat diagnoses after the first new diagnosis. | Age, sex, ART status, calendar year. |
| % HIV positive tests among all tests ("positivity") | Age, sex, calendar year. |
| % of new HIV diagnoses among all tests ("yield of new diagnoses") | Age, sex, calendar year. |
| **Testing history** | |
| Proportion of ≥15 years old ever tested for HIV | Age, sex, PLHIV status, calendar year. |
| Proportion of ≥15 years old tested for HIV in the past year | Age, sex, PLHIV status, calendar year. |
| **Knowledge of HIV status among PLHIV** | |
| Proportion of PLHIV who know their status ("first 90 estimates") | Age, sex, calendar year. |

ART: antiretroviral therapy; PLHIV: people living with HIV.

**Table S5.** Posterior estimates of the main model parameters for Côte d'Ivoire, Malawi, and Mozambique (full data calibration).

| **Parameters** | **Symbols** | **Côte d'Ivoire (95%CrI)** | **Malawi (95%CrI)** | **Mozambique (95%CrI)** |
|---|---|---|---|---|
| RR testing for men in 2005 | $M_1(2005)$ | 0.69 (0.53-0.84) | 0.73 (0.65-0.82) | 0.41 (0.35-0.49) |
| RR testing for men in 2012 | $M_1(2012)$ | 0.38 (0.34-0.43) | 0.49 (0.46-0.52) | 0.37 (0.32-0.42) |
| RR re-testing in 2010 | $RR_{Test}(2010)$ | 3.49 (2.84-4.22) | 1.09 (1.04-1.16) | 7.10 (6.12-7.61) |
| RR re-testing in 2015 | $RR_{Test}(2015)$ | 3.74 (3.21-4.31) | 1.20 (1.09-1.40) | 7.22 (6.39-7.65) |
| RR for PLHIV testing unaware | $RR_{Unaware}$ | 1.32 (0.93-1.62) | 1.27 (1.14-1.40) | 1.33 (1.11-1.51) |
| RR re-testing PLHIV aware | $RR_{Aware}(2010)$ | 4.82 (3.21-6.19) | 1.30 (0.96-1.73) | 3.75 (1.88-5.75) |
| RR re-testing PLHIV aware | $RR_{Aware}(2017)$ | 2.41 (1.37-3.80) | 1.29 (0.69-2.26) | 3.85 (1.86-5.91) |
| RR re-testing PLHIV on ART | $RR_{ART}$ | 0.73 (0.26-0.95) | 0.09 (0.01-0.51) | 0.24 (0.02-0.82) |
| RR for men aged 25-34 years | $A_{1,2}$ | 1.37 (1.14-1.64) | 1.41 (1.29-1.56) | 1.16 (0.97-1.38) |
| RR for men aged 35-49 years | $A_{1,3}$ | 0.90 (0.74-1.08) | 0.77 (0.70-0.84) | 0.62 (0.50-0.77) |
| RR for women aged 25-34 years | $A_{2,2}$ | 0.85 (0.74-0.97) | 1.10 (1.04-1.17) | 0.83 (0.75-0.93) |
| RR for women aged 35-49 years | $A_{2,3}$ | 0.89 (0.79-1.00) | 0.64 (0.60-0.67) | 0.37 (0.33-0.42) |
| Factor for % of OI tested for HIV | $\kappa$ | 0.60 (0.30- 1.40) | 1.10 (0.30-1.70) | 1.30 (0.50-1.70) |

95%CrI: 95% credible intervals; ART: antiretroviral therapy; OI: opportunistic infections; PLHIV: people living with HIV; RR: rate ratio.
